# Rational inattention in neural coding for economic choice

**DOI:** 10.1101/2024.09.20.614193

**Authors:** Justin M. Fine, Rubén Moreno-Bote, Benjamin Y. Hayden

## Abstract

Mental operations like computing the value of an option are computationally expensive. Even before we evaluate options, we must decide how much attentional effort to invest in the evaluation process. More precise evaluation will improve choice accuracy, and thus reward yield, but the gain may not justify the cost. *Rational Inattention theories* provide an accounting of the internal economics of attentionally effortful economic decisions. To understand this process, we examined choices and neural activity in several brain regions in six macaques making risky choices. We extended the rational inattention framework to incorporate the foraging theoretic understanding of local environmental richness or reward rate, which we predict will promote attentional effort. Consistent with this idea, we found local reward rate positively predicted choice accuracy. Supporting the hypothesis that this effect reflects variations in attentional effort, richer contexts were associated with increased baseline and evoked pupil size. Neural populations likewise showed systematic baseline coding of reward rate context. During increased reward rate contexts, ventral striatum and orbitofrontal cortex showed both an increase in value decodability and a rotation in the population geometries for value. This confluence of these results suggests a mechanism of attentional effort that operates by controlling gain through using partially distinct population codes for value. Additionally, increased reward rate accelerated value code dynamics, which have been linked to improved signal-to-noise. These results extend the theory of rational inattention to static and stationary contexts and align theories of rational inattention with specific costly, neural processes.

## INTRODUCTION

Why do we “pay” attention? Acquiring information through attention requires effort, which is costly (Botvinick and Braver, 2015; Shenhav et al., 2017; Stigler, 1961). The decision to pay attention should be made just like any other cost-benefit decision: by estimating its cost and comparing it with the net benefit expected from its expenditure. This cost-benefit logic applies to any process that requires attention, including evaluating options in choice. Evaluation is an attentionally demanding computational sampling process (Bakkour et al., 2019; Krajbich et al., 2010; Rangel et al., 2008; Rich and Wallis, 2016), and is, therefore, cognitively costly (Lieder and Griffiths, 2019). Rational choosers, then, should exert more attentional effort in evaluation when it is valuable to do so (Enke, 2024; Glimcher, 2022; Polania et al., 2024).

Conversely, when the benefits of evaluation are reduced, rational choosers should withdraw attention and rely on approximation, even at the risk of choice errors. This is the core logic of the *rational inattention theory*, which formalizes the economic principles by which we allocate our attentional effort (Dean and Neligh, 2023; Gabaix et al. 2019; Gershman and Burke, 2022; Matêjka et al., 2015; Sims, 2003; Woodford, 2009). Behavioral studies have provided evidence in favor of the predictions of rational inattention by showing how changes to available rewards can modulate intertemporal choice precision (Gershman and Bhui, 2020), risky choice (Dean and Neligh, 2023), and alter perceptual discrimination learning (Grujic et al., 2022).

From the rational inattention perspective, willingness to expend attentional effort should be motivated by available reward. In some cases, the value of attending might be determined by the learned average reward of an environment (Mikhael et al., 2021). However, in other cases, the determiners of attentional effort can be more complex. Most environments exhibit fluctuating richness levels around an average reward. And decision-makers need to predict in advance whether the future attentional effort will pay off. Foraging theory tells us that decision-makers can predict the value of this future attentional effort by monitoring the local richness or local reward rate in comparison to the global average reward, and tune their strategy to its variation (Charnov, 1976; Hayden, 2018; Stephens and Krebs, 1986). Specifically, in richer contexts, foragers should invest more effort because the effort leads to greater and sooner expected payoff, and vice versa (Shadmehr and Ahmed, 2020). Indeed, there is evidence that even in static environments with a stable average reward, decision-makers’ efforts (e.g., vigor) in choice can be motivated by the recent reward rate in accordance with foraging principles (Yoon et al., 2018).

Thus, an increase in local reward rate is likely then interpreted as increased environmental richness, and would therefore promote attentional effort in offer value evaluation. Thus, a foraging perspective offers a principled expansion of what motivates rationally inattentive behavior from a dynamic environment – which has known changes in rewards – to the broader case where reward rates have to be calculated and environmental reward statistics are not fully known.

While rational inattention offers a powerful explanation for behavioral data, the neural processes that support its implementation remain unknown. Here, we examined a large dataset of behavior and neural activity in six rhesus macaques performing a risky choice task (Strait et al., 2014). We found, confirming predictions made by our extension of rational inattention theory, choice accuracy is positively correlated with recent reward rate. Furthermore, baseline and evoked pupil are both higher in richer reward contexts, supporting the idea that these improvements are due (at least in part) to attentional effort. We examined responses of single neurons in the ventral striatum, orbitofrontal cortex, pregenual cingulate cortex, and posterior cingulate cortex. In all regions, we observed a systematic decodability of reward rate before the start of the trial. In VS and OFC, increases in reward rate resulted in improved value decodability during offer evaluation; this increase could be directly attributed to an increase in neural gain.

Reward rate also partitioned the neural geometries for value coding into semi-orthogonal subspaces, while value was still decodable in both subspaces. This result deviates from a pure neural gain model (McAdams and Maunsell, 1999; Salinas and Thier, 2000) in which tuning to value would be fixed across reward rate contexts, and subsequently would predict aligned value subspaces. Instead, the semi-orthogonalization of subspaces supports an abstract value code that is bound to different reward rate contexts (subspaces) with different gains (Johnston and Fine; 2024; Bernardi et al., 2020); this points to gain control operating as a distributed population code rather than amplitude modulation of neurons with a fixed tuning to value (Xie et al., 2022). In other words, partially distinct population codes were used for different gains. Finally, we found the value coding subspaces during evaluation were dynamic rather than persistently stable (Enel et al., 2020; Goldman, 2009; Stokes et al., 2013), and, specifically, that richer contexts led to faster changing (more dynamic) codes. These neural results therefore link rationally inattentive behavior with specific, likely costly, neural processes, and show how attentional effort operates by changing population ensemble codes.

## RESULTS

We analyzed choices made by six rhesus macaques performing a two-option **asynchronous gambling task** (Strait et al., 2014; Fine et al., 2023; **Figure 1A**). On each trial, two offers appear in sequence (one-second asynchrony) on opposite sides of a computer screen (left or right). Then the two offers reappear, simultaneously, and the subject makes a choice by shifting gaze and fixating their preferred offer. Each offer is defined by a probability (0-100%, 1% increments) and stakes (large or medium reward, 0.240 and 0.165 mL juice, respectively). The probabilities and stakes associated with both offers are randomized for each option. The order of presentation (left first vs. right first) is randomized by trial.

**Figure 1.**
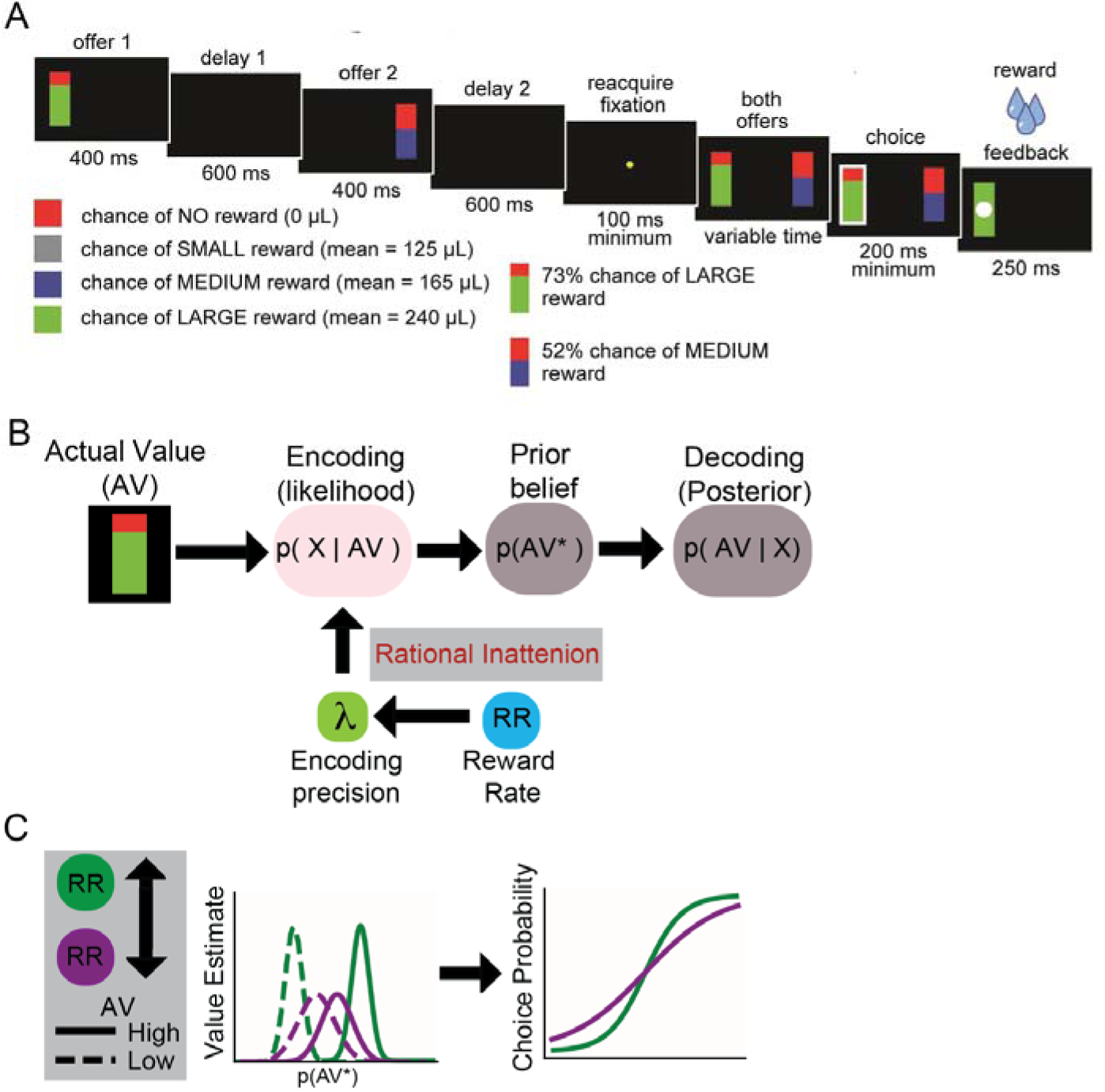
Task outline, description of rational inattention theory, and predictions. **A.** asynchronous gambling task. On each trial, subjects first sees an offer (risky option) on either the left or right. Following a 600 ms blank delay, a second offer appears for 400 ms; after another 600 ms delay, both options reappear, subjection chooses, and reward is given. **B.** Rational inattention is a Bayesian observer theory that describes how the actual values (AV) on offer are encoded as a noisy percept (likelihood) and combined with a prior belief about the distribution of values (p(AV*)). Rational inattention theory proposed the likelihood precision is enhance by a larger reward expectation. A foraging perspective on Rational inattention theory posits that the precision of the likelihood is modulated by the local reward rate. **C.** As a consequence, in the risky choice task, when reward rate is high (green distributions), the internally estimated offer value deviates more from the mean of the prior distributions of values (p(AV*)) and tend towards the correct value. While in low reward rate contexts, estimated values exhibit a regression to the prior mean. Neurally, this means value decodability between low and high offer values should be more accurate in high versus low reward rate conditions. In addition, the choices should become more accurate as reward rate increases and should be reflected as a steepening of a choice logistic curve.

Because a subject’s perception of the offer value is uncertain, it must infer the actual offer value (**Figure 1B**). Bayesian observer models describe how an optimal observer can combine their noisy perception of a stimulus (likelihood) with a prior belief over its values (Ma et al., 2023). Typical Bayesian models assume the likelihood noise is extrinsic and not under the observer’s control. Rational inattention generalizes this idea by acknowledging that observers can expend attentional effort to improve perception (Sims, 2003; Woodford, 2009). However, optimizing observers should only invest this effort when the expected benefits outweigh the costs. In our task, a reward rate proxy is assumed to motivate or set attentional effort or coding precision (**Figure 1B**; Mikhael et al., 2021). Therefore, increases in reward rate should yield more precise value estimates and, therefore, more optimal choices when choosing between the values because the options are more discriminable (**Figure 1C**).

### Subjects make better choices when reward context is richer

We predicted subjects would devote more attention, and thus show more accurate choices, when the local reward rate was higher. Based on our earlier work relating choice strategy to recent outcomes, we defined reward rate as an exponentially decaying function over recent rewards (in this case, 3 trials) compared to the subject’s global reward rate (across sessions, cf. Hayden et al., 2008 and 2011). To quantify changes in choice accuracy, we performed a logistic model of subject choices using regressors (1) for difference in expected value between the two offers (Δ*EV*), (2) reward rate, and (3) the interaction of reward rate with Δ*EV* variables. All six subjects exhibited a higher model conditional probability (AIC weights all > 0.62 for each subject; **Figure 2A**) in favor of a model with Δ*EV* and a reward rate interaction of Δ*EV.* In general, Δ*EV* scaled positively with reward rate (**Figure 2B**), meaning subjects were more accurate at discriminating close stimuli when recent reward rate was greater (**Figure 2C**; all subjects: t(320)=14.36, *p* < 0.00001).

**Figure 2.**
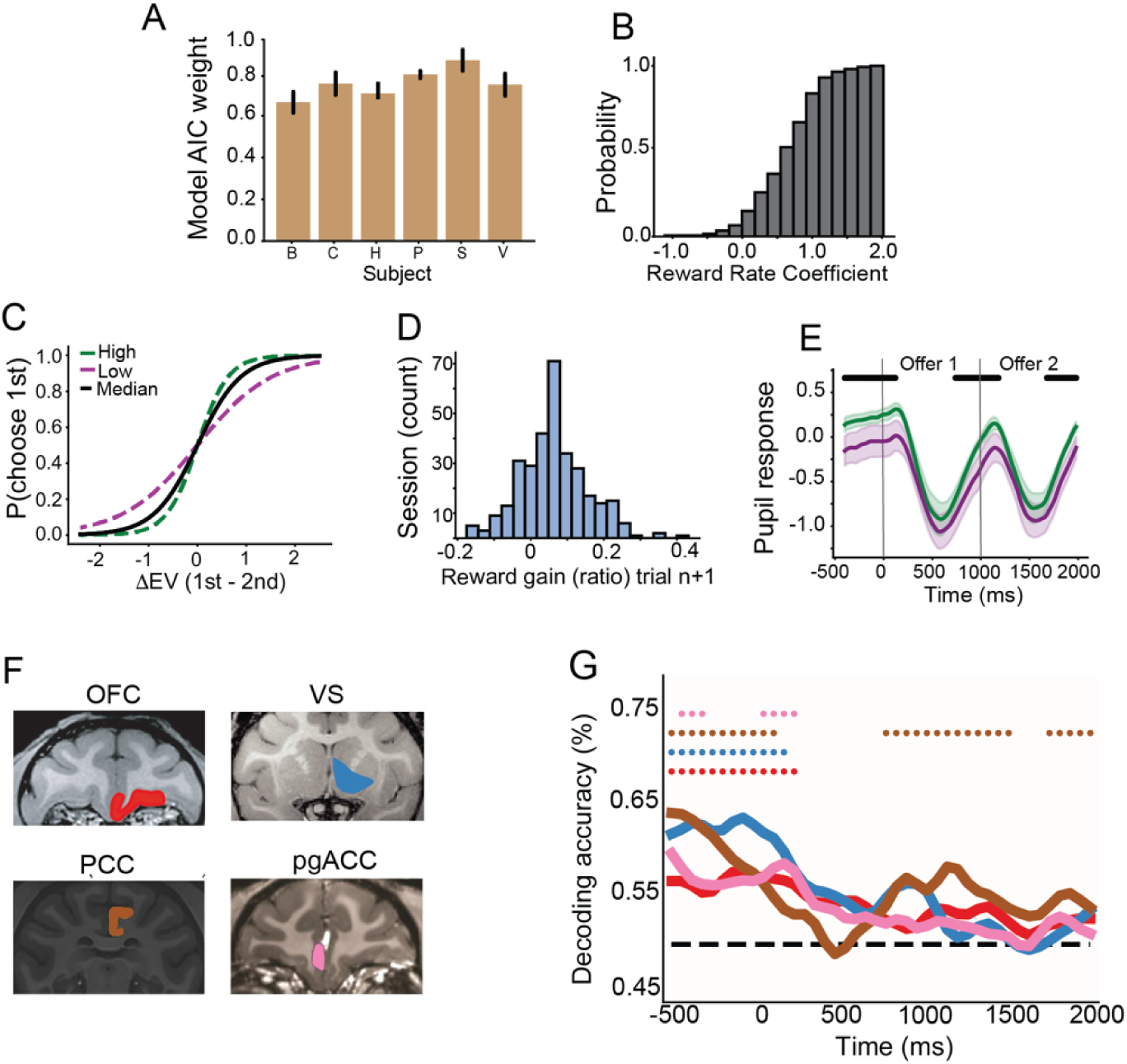
Logistic choice model, pupillometry, brain areas and reward rate decoding. **A.** AIC model weights, showing probability in favor of the logistic choice model with a reward rate x Δ*EV* interaction term. Weights are the mean and standard error across for each subject, taken over each subject’s sessions. The weights were all greater than 0.5, indicating this model was favored for all subjects. **B.** The cumulative density choice function of the reward rate x value interaction regression coefficients for all session logistic models. **C.** Logistic choice curve from average model coefficients fitted across six subjects. Curves are shown for the low reward rate context (purple), high reward rate context (green) and the median reward rate (black line). Subjects show more optimal choices (steeper slope) in the high reward rate condition. **D.** Average reward gain defined as the normalized ratio of high reward rate to low reward rate contexts. Reward gain increases for the n+1 trial defined after the 3 trial reward rate window. Subjects gained more reward on these trials following high reward rates. **E.** Baseline corrected mean pupil across subjects, for both the low (purple) and high (green) reward rate conditions. Note that values are greater even before an offer appears. Shading: standard error. Black dots: time points with significant differences. **F.** MRI coronal slices showing the 4 different core reward regions that were analyzed. **G.** The decoding of reward rate across all brain areas (lines) and their significant points (dots) for each brain area. Each brain area is colored according to **E.** OFC: red; VS: blue; PCC: brown; pgACC: pink.

The extra effort applied had tangible results. We computed the reward gain using trials that were non-overlapping from those used to compute the reward-rates (ie., the next trial). Specifically, we calculated the normalized ratio of subsequent reward gained on high versus low reward rate. On average, higher reward rate resulted in a gain of approximately 10% of the global average reward size (**Figure 2D**; specifically, an additional 0.24 mL of juice; t(320) = 12.61, *p* < 0.00001) per trial. This result places a specific value of attentional effort in units of juice volume - the relevant unit for these monkeys in this context. This result implies that, *ceteris paribus*, the subjective, internal, cost of applying attentional effect in the amount allocated in the higher reward rate context was equivalent to 0.24 mL of juice. These results accord with the prediction that higher reward rates may incentivize more attention to option values, leading to more optimal choices (**Figure 2D**).

### Pupil responses reflect increased reward rate context

We hypothesized that these changes in behavior reflect changes in the allocation of attentional effort. To obtain complementary evidence in favor of the attention hypothesis, we examined pupil size. Pupil size has long been considered a measure of attentional effort and overall arousal state more generally (Strauch et al., 2022; Urai et al., 2017; van der Wel and van Steenbergen, 2018), and is also linked to neural gain (Aston-Jones and Cohen, 2005; Eldar et al., 2013). For this reason, tonic pupil differences have been used to index rationally inattentive behavior in mice (Grujic et al., 2022).

To determine whether these patterns of pupil activity apply in our macaques, we acquired pupil size measures in three of our subjects (V, S, and P). In all three, pupil size increased steadily between fixation onset and the first offer window (**Figure 2E**). Overall pupil size was locked to key events across the offer epochs and their delay periods. Using a permutation t-test (false discovery rate corrected *p* < 0.05) that split pupil responses on high vs. low reward rate, we found several contiguous points where pupil response was larger for a high reward rate context (**Figure 2E**). Specifically, we found higher pupil responses for high reward rate in the baseline period (∼ −400:0 ms, where 0 indicates offer 1 onset). We also found larger pupil sizes in both offer evaluation periods (∼ 0:200 ms and 1000 - 1200 ms) and during both memory periods for both offers (∼600:1000 ms and 1600:2000 ms). These results support the hypothesis that increases in reward rate are likely paired with increased attentional effort, both before and during evaluation.

### Pre-trial neural activity encodes local reward rate context

We next examined the dependence of neural activity on local reward rate in four brain regions, orbitofrontal cortex (OFC), pregenual anterior cingulate cortex (pgACC), posterior cingulate cortex (PCC), and ventral striatum (VS). To increase statistical power, we combined neurons from central OFC (cOFC, Areas 13 and 11) with those in medial OFC (mOFC, Area 14o, Ongur and Price, 2000). We refer to this larger area as OFC in results. Together, these four regions (**Figure 2F**) constitute the majority of the core reward network, a set of regions whose neural activity robustly encodes values of offers and outcomes (Barta et al., 2013; Clithero et al., 2014).

We first asked whether we could decode reward rate (low vs. high, median-split) in each region. To do this, we used a linear support vector machine (SVM, **Methods**). Reward rate was decodable during the baseline period (−500 to 0 ms) in all four brain regions (**Figure 2G**, permutation test, *p* <0.05, false discovery rate corrected). Reward rate information only became decodable again in PCC at the onset of the second offer and was maintained throughout the second offer window. Thus, all four core value regions contain a neural signal for reward rate context that presumably covaried with an attentional allocation process. Because the reward rate decoding was predominantly found during the baseline, attentional allocation was likely set before offers were evaluated.

### Value information is gain modulated by reward rate context

If subjects operated commensurate with predictions from rational inattention theory, then attentional effort should increase coding fidelity during offer evaluation. Attention alters gain in single neurons (David et al., 2008; Hermann et al., 2010; McAdams and Maunsell, 1999; Treue and Maunsell, 1996). We therefore predicted that larger reward rate contexts would show gain-enhanced value coding relative to low-reward trials. Notably, gain changes have a direct translation to population coding: it is established that increases in overall tuning gain to a variable – value here – directly translate to increased distance between stimulus representations in their neural geometry (Kriegeskorte and Wei, 2021; Johnston and Fine, 2024). Larger distances between neural population codes predict higher decoding accuracy. Therefore, we can assess the prediction of population gain change by looking directly at change in neural value decoding for different levels of reward rate. We used linear SVMs to classify value (low vs high, median split), separately for each reward rate context. We can visualize the population activity in value space by projecting the mean firing activity onto the decoding hyperplanes. Examining these projections indeed suggests OFC and VS have a higher gain for value coding during high reward rate contexts (**Figure 3A-D**). Next, we provide quantitative evidence for this observation.

**Figure 3.**
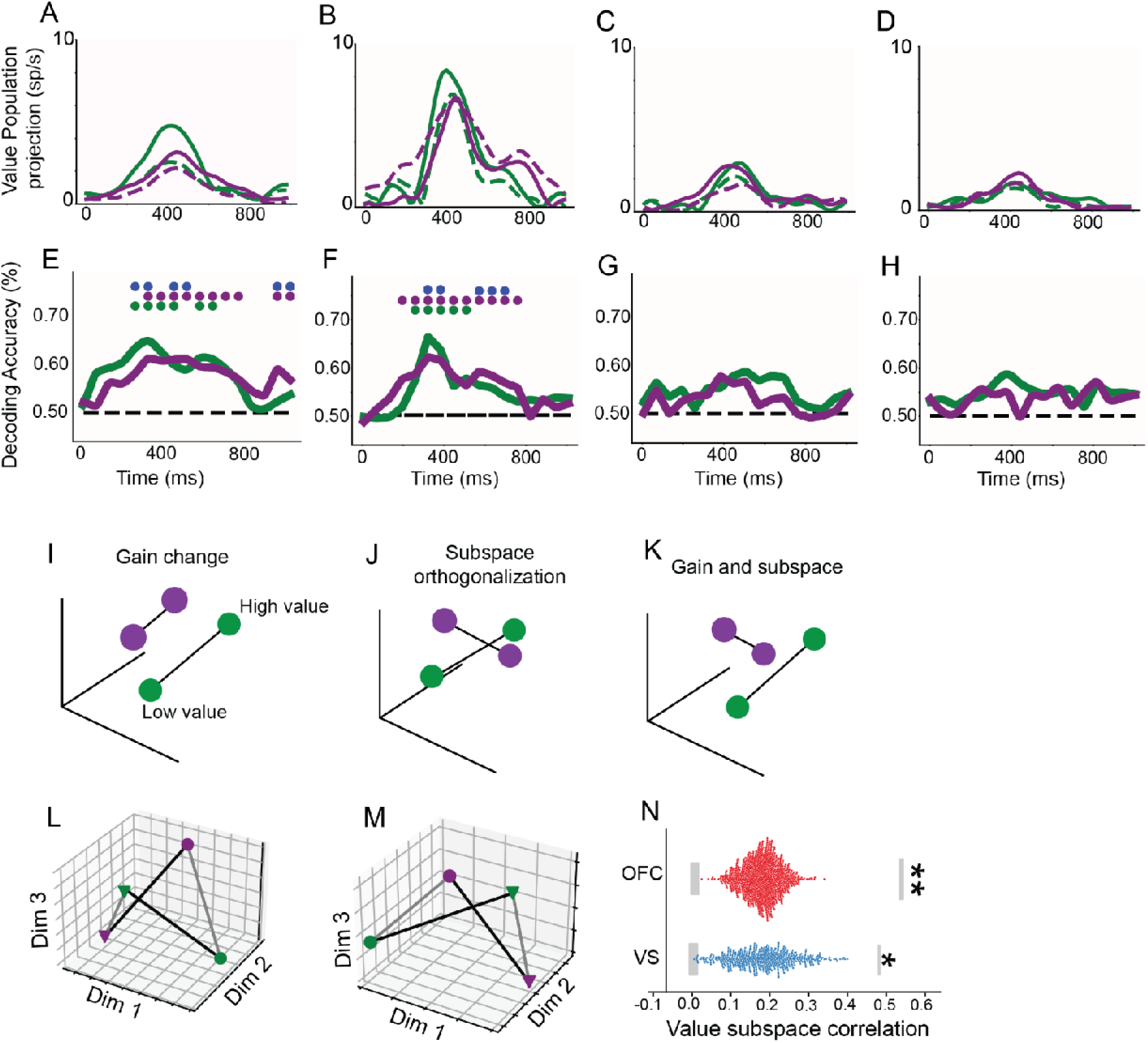
Value subspace projection, Value decoding, Neural geometries, Value Subspace Correlations. **A-D** (OFC, VS, PCC, pgACC). Projection of each brain area’s firing rate activity onto the value decoding subspaces– showing the population firing rate in the value space. Purple and green lines are the low and high reward rate projections, respectively. Dashed and solid lines are low and high value conditions, respectively. This shows how gain changes appear in the population coding of value; for example, in **A.** and **B.** the difference in solid and dashed green lines (high reward rate) is evidenced by greater than that for solid and dashed purple (low reward rate) conditions. **E-H** Value decoding across both reward rate conditions and their difference. Lines are the mean decoding accuracy. Accuracy for low and high reward rate conditions are purple and green, respectively. Significant decoding is shown as a matching color dot. Blue dots show significant differences in decoding accuracy between reward rate conditions. **E.** OFC. **F.** VS. **G.** PCC. **H.** pgACC. **I-K.** Different three-dimensional neural population geometries corresponding to different hypothetical population coding scenarios for increased reward rate driving coding. **I.** Pure gain coding models would yield aligned differences in decoding accuracy between conditions, with greater distance between points as decoding accuracy increases (purple: low reward rate; green: high reward rate). Values subspaces defined by decoder hyperplanes (lines connecting points) would also be parallel (non-orthogonal). **J.** The subspaces can also be rotated and imperfectly aligned (orthogonality). More orthogonal subspaces are unaligned with pure gain models, as this implies neuron value tuning is not invariant to auxiliary conditions (here, reward rate). **K.** An example of a neural geometry that exhibits both gain coding and subspace rotation. **L.** Neural value geometry (subspaces) for OFC estimated with multidimensional scaling. **M.** same as **L.** for VS neural geometry. **N.** swarmplots showing bootstrap distributions of subspace correlations between low and high reward rate contexts. Gray box around zero correlation shows noise level for each area (*p<0.05, **p<0.01), and upper gray box shows ceiling for orthogonality criterion.

Our analysis focused on the first offer window (0-1000 ms) as it is separated from any processes involved in option comparisons that happen in the second offer window (Yoo and Hayden, 2020). In both OFC and VS, value was decodable during the high reward rate and low-reward rate conditions (**Figure 3E-F**). Value was not decodable in any of the other regions during this time-window (**Figure 3G-H**).

Next, we tested the hypothesis that population gain is greater in high reward rate contexts. We compared decoding accuracies between reward rate levels using Wilcoxon-rank tests (permutation and false discovery rate corrected *p* < 0.05). During the offer evaluation period (0-400 ms), we observed this gain coding difference in both OFC and VS (**Figure 3E-F**). These results indicate OFC and VS carry a change in evaluation that aligns with the rational inattention predictions. Surprisingly, during the memory window (the subsequent 600 ms during which the monitor was blank) we observed the opposite direction of differential decoding in both OFC and VS – low reward rate trials exhibited a higher accuracy; nonetheless, it is notable this latter window likely involves distinct working memory processes rather than the evaluation process that requires effortful attention and gain control. Thus, we provide evidence for population gain effects for online evaluation in OFC and VS; for this reason, we do not further evaluate pgACC and PCC regions in this study.

### Value coding subspaces are semi-orthogonal between reward rate contexts

We next asked how reward rate context changes the geometry of value coding. Geometry changes could occur instead of or in conjunction with a gain change (**Figure 3I-K**). In the language of vector spaces, attentional effort could alter the vector length (gain; **Figure 3I**) or the angle (geometry; **Figure 3J**), or both at the same time (**Figure 3K**). A pure gain model would predict highly similar value subspaces between reward rate contexts; this is because pure gain control would result from neurons that are invariantly tuned to value. In contrast, a population code could still exhibit attentionally based gain control even if the single neuron tuning to value shifts across reward rate contexts, predicting semi-orthogonal subspaces. To distinguish these two hypotheses, we quantified the alignment between value coding subspaces – defined by the decoder hyperplanes – in high- and low-reward rate conditions (Libby and Buschman, 2021). Specifically, we took the decoder weights to instantiate the linear subspace for value separately for each reward rate context. We estimated alignment (that is, orthogonality) by correlating the SVM decoder weights, effectively performing targeted dimensionality reduction (Kimmel et al., 2020). To avoid neural confounds of value comparison and choice, we focused on an offer evaluation window in which decoding of value was highest (100-400 ms; **Figure 3E-F**).

We found in both OFC (**Figure 3L**) and VS (**Figure 3M**) that subspace correlations between low and high reward rate were semi-orthogonal (**Figure 3N**). Specifically, responses in both regions were lower than the noise ceiling (p <0.0001, see Methods) and greater than a shuffle-based noise floor (OFC: p = 0.002; VS: p = 0.038; **Figure 3N**). This result indicates that the gain differences (reward rate context) in value decoding (in this window) are not a simple modulation of neurons with fixed tuning to value. This is because maintaining a decodable value signal while employing semi-orthogonal subspaces requires that some neurons change their tuning to value across reward rate contexts. Thus, these results suggest gain control over value operates by partitioning population codes based on whether they were evaluated with either low or high attentional effort (reward rate).

### Temporal dynamics of value coding subspaces are modulated by reward rate context

Neural codes are often dynamic; the type of dynamics they exhibit can be indicative of different underlying network schemata (Murray et al., 2016; Stroud and Lengyel, 2024; Wang et al., 2023). Previous modeling work has shown dynamic codes are largely driven by networks with effectively feedforward connectivity (Goldman, 2009; Stroud and Lengyel, 2024). The same modeling has shown these dynamic, feedforward codes can exert an information processing advantage compared to classic linear integrators/attractors: the increased dynamicism of effective feedforward networks may amplify the signal to noise ratio of processed inputs (Ganguli et al., 2008; Goldman, 2009; Hennequin et al., 2012; Murphy and Miller, 2009). We hypothesized such network-based signal amplification could support attentional effort control through code dynamics. Thus, to quantify the dynamics of value coding in OFC and VS, we asked whether the neural decoding value subspaces are stable or dynamic, and in the dynamic case, how its dynamics are affected by reward rate context. We used cross-temporal decoding (CTD) of value with a linear SVM decoder (Stokes et al., 2013; Meyers et al., 2008). In CTD, a decoder is trained on one time window and tested for generalization on another (**Methods**). Thus, CTD tests how well a value code at one time-point generalizes to another time-point. High CTD throughout the offer window implies a stable code.

In both OFC and VS, and in both high and low reward rate contexts, we found significant temporal generalization (**Figure 4A-B**, OFC, and **Figure 4D-E**, VS). For example, both regions exhibited generalization across the offer evaluation window (0-400 ms; **Figure 4A-B**, OFC, and **Figure 4D-E**, VS). Both regions also exhibited some generalization with the delay window (400-1000 ms; **Figure 4A-B**, OFC, and **Figure 4D-E**, VS). However, this generalization was relatively short-lived in both windows: the coding subspaces for evaluation and memory periods were distinct and did not generalize to one another. These results accord with previous studies indicating the subspaces for evaluation or online perception are rotated into a distinct subspace during memory (Libby and Buschman, 2021; Johnston and Fine 2024; Yoo and Hayden, 2020; Tang et al., 2020).

**Figure 4.**
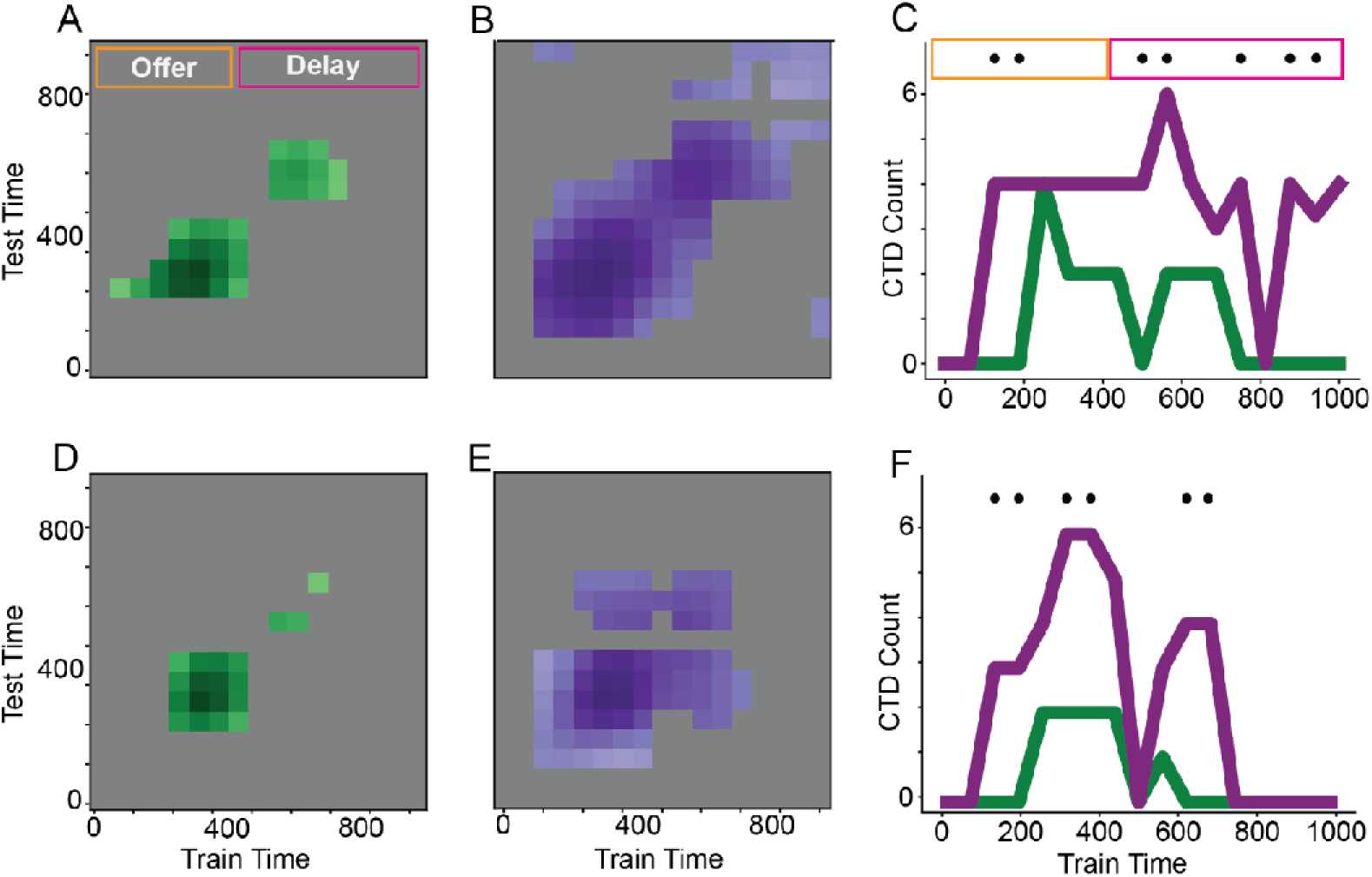
Cross-temporal decoding (CTD). **A - B** CTD across the first offer window for OFC in the high reward rate (**A**; green) and low reward rate (**B**; purple) contexts. Training-time points for the SVM are on the x-axis and test-time points are on the y-axis. Significant points of CTD are colored in. Both plots show substantially more CTD in low reward rate contexts (**B)**, as verified in **C.** showing the counted number of significant time points of CTD for OFC, for each training time point. Black dots indicate significantly different counts between reward rate contexts; all significant points indicated low reward rate contexts (purple) had a higher CTD (more stable) compared to high reward rates (green; more dynamic). **D-E**. shows the CTD for VS region, with **D.** showing high reward rate CTD and **E.** showing low reward rate contexts. **F.** shows a pattern of CTD counts (dynamics) where low reward rate contexts where more stable than high reward rate contexts.

Next, we quantified the differences in value subspace dynamics between each reward rate context by comparing the proportion of significant off-diagonal terms (**Figure 4C**, OFC and **Figure 4F**, VS). Across windows and both OFC and VS, we found a more dynamic code for high-reward rate contexts (false discovery rate, all significant points *p* < 0.05). Put differently, low-reward contexts exhibited more temporal generalization (stability) in value codes across time. These results are consistent with the hypothesis that the resulting reward rate differences in value code dynamicism reflect an amplification of value coding precision.

## DISCUSSION

We used the theory of rational inattention (RI) to understand choice behavior and reward-related neural responses in macaques performing a risky choice task. We find, consistently across six subjects, that behavioral performance, as estimated by choice variability, waxes and wanes across trials. Our central behavioral result is that this variability is not random but is enhanced in temporally local rich contexts (e.g., a pattern of recent wins). This pattern is predicted by an extension of rational inattention theory that incorporates foraging theory, in which evaluation requires costly allocation of attention and the allocation of attention is determined by temporally local reward rate. Specifically, it predicts that decision-makers increase the precision of evaluation in richer reward rate contexts. The hypothesized link between evaluation and attention is further supported by our complementary pupil size results. Together, these results provide strong behavioral and neural evidence in favor of the principles of rational inattention as a basis for controlling attentional effort in option evaluation.

One important feature of our study is extending rational inattention principles from shifting reward contexts to include static (including stationary) contexts. Previous studies have generally focused on contexts in which the optimal allocation of attention covaries with the dynamic variability in payoff structure within that context (Gershman and Burke, 2022; Grujic et al., 2022). In contrast, we propose that attentional allocation decisions are based on internal estimates of cost-benefit, which can be driven by stochastic variability in a stationary environment. This idea can then be rationalized with the foraging theoretic idea that local environmental richness (reward rate) should motivate investment (Charnov, 1976; Hayden et al., 2011; Yoon et al., 2018). Our approach, then, extends the ideas elaborated in previous studies of rational inattention and models (Grujic et al., 2022; Gershman and Burke, 2022). Specifically, an insightful theory was recently developed that connected the motivating signal of rationally inattentive perceptual control to average-rewards and tonic dopamine (Mikhael and Gershman, 2021).

Our neural results provide a potential neural basis for the costs and benefits of attentional effort. During trials with greater attentional allocation, value responses in OFC and VS are enhanced with a gain modulation, and as a result, value decodability increases. This effect supports the assertion that attentional effort is costly because it requires more metabolically costly spikes (Laughlin et al., 1998). This cost was presumably counterbalanced by harvesting of additional juice reward; indeed, we show that subjects gained more reward on those trials in which gain was higher. However, this was not the only effect of attentional effort; attention also systematically alters population coding geometries in a way that deviates from a pure gain model of attention. Specifically, we found semi-orthogonal subspaces for value coding in both OFC and VS between the different reward rate contexts. One possible explanation for the subspace distinction is that it reflects the extent to which a value code in a reward rate context is projected from an encoding-oriented subspace to a comparison-oriented or choice-oriented subspace (Elsayed et al., 2016; McGinty and Lupkin, 2023; Panichello and Buschman, 2021;Yoo and Hayden, 2020).

Why would attentional effect have this effect? We speculate that these distinct subspaces may bind the value code to different reward rate contexts that convey an evaluation confidence-like signal (Pouget et al., 2016). Consider, for example, that in low reward rate (and thus low-attention) contexts, subjects may only weakly sample the stimulus. By using partially distinct subspaces that are tagged with a confidence signal, this weak sampling of value can be translated to downstream neurons involved in choice comparisons, allowing them to discriminate whether the encoded value was done under low-or high-attention. Coding of confidence signals would be consistent with previous work showing OFC subregions can code for subjective value (Padoa-Schioppa, 2011) and confidence signals (De Martino et al., 2013; Gherman and Phillistades, 2018; Lebreton et al., 2015). More generally, finding semi-orthogonal value subspaces indicates the value code strikes a balance between being able to bind the encoded value with the reward rate context, while also being able to generalize the value code between contexts (Barak et al., 2013; Bernardi et al., 2020; Nogueira et al., 2023; Johnston and Fine, 2024).

Our finding that richer reward rate contexts produce more dynamic value codes is important for understanding how attentional effort controls value coding accuracy. We speculate on the computational benefits of using dynamic codes rather than stable attractors and drift-diffusion models found in common models of evaluation and choice (Hunt et al., 2012; Krabich et al., 2010; Rustichini and Padoa-Schioppa, 2015). We hypothesize that dynamic codes allow an amplification of the inputs used to evaluate the offer value, improving coding fidelity. This idea is supported by several modeling studies showing that dynamic codes are both driven by networks with an effectively feedforward connectivity structure (non-normal network) and have the benefit of amplifying the signal to noise ratio of processed inputs (Baggio et al., 2020; Stroud and Lengyel, 2024). The reason for this amplification is because when an input is processed in a feedforward chain, and it’s projected earlier into that chain compared to later, it has more chances to transiently amplify that signal by processing through more connections and mitigating the impact of noise (Goldman, 2009). Importantly, if such a feedforward process supports offer evaluation, then making a code more dynamic by processing it through more steps, then the evaluated offer signal fidelity will be amplified. This fact in turn points to another link between our physiological findings and the benefits of attentional control: more attentional effort may convey more accurate value information via dynamic coding.

## Acknowledgements

We thank Tommy Blanchard, Tyler Cash-Padgett, Marc Mancarella, Brianna Sleezer, Caleb Strait, Maya Wang, and Michael Yoo for assistance with data collection.

## Methods

### Surgical procedures

All procedures were approved by either the University Committee on Animal Resources at the University of Rochester or the IACUC at the University of Minnesota. Animal procedures were also designed and conducted in compliance with the Public Health Service’s Guide for the Care and Use of Animals. All surgery was performed under anesthesia. Male rhesus macaques (*Macaca mulatta*) served as subjects. A small prosthesis was used to maintain stability. Animals were habituated to laboratory conditions and then trained to perform oculomotor tasks for liquid rewards. We placed a Cilux recording chamber (Crist Instruments) over the area of interest. We verified positioning by magnetic resonance imaging with the aid of a Brainsight system (Rogue Research). Animals received appropriate analgesics and antibiotics after all procedures. Throughout both behavioral and physiological recording sessions, we kept the chamber clean with regular antibiotic washes and sealed them with sterile caps.

### Recording sites

We approached our brain regions through standard recording grids (Crist Instruments) guided by a micromanipulator (NAN Instruments). All recording sites were selected based on the boundaries given in the Paxinos atlas (Paxinos et al., 2008). In all cases we sampled evenly across the regions. Neuronal recordings in OFC were collected from subjects P and S; recordings in rOFC were collected from subjects V and P; recordings in vmPFC were collected from subjects B and H; recordings in pgACC were collected from subject B and V; recordings from PCC were collected from subject P and S; and recording in VS were collected from subject B and C.

We defined c**OFC** as lying within the coronal planes situated between 28.65 and 42.15 mm rostral to the interaural plane, the horizontal planes situated between 3 and 9.5 mm from the brain’s ventral surface, and the sagittal planes between 3 and 14 mm from the medial wall. The coordinates correspond to both areas 11 and 13 in Paxinos et al. (2008). We used the same criteria in a different dataset (Blanchard et al., 2015).

We defined **mOFC 14o** as lying within the coronal planes situated between 29 and 44 mm rostral to the interaural plane, the horizontal planes situated between 0 and 9 mm from the brain’s ventral surface, and the sagittal planes between 0 and 8 mm from the medial wall. These coordinates correspond to area 14m in Paxinos et al. (2008). This dataset was used in Strait et al., 2014 and 2016, and corresponds to the same region used in Jurewicz et al., (2024) and Maisson et al. (2021).

We defined **pgACC 32** as lying within the coronal planes situated between 30.90 and 40.10 mm rostral to the interaural plane, the horizontal planes situated between 7.30 and 15.50 mm from the brain’s dorsal surface, and the sagittal planes between 0 and 4.5 mm from the medial wall. Our recordings were made from central regions within these zones, which correspond to area 32 in Paxinos et al. (2008). Note that the term 32 is sometimes used more broadly than we use it, and in those studies encompasses the dorsal anterior cingulate cortex; we believe that that region, which is not studied here, should be called area 24 (Heilbronner and Hayden, 2016).

We defined **PCC 29/31** as lying within the coronal planes situated between 2.88 mm caudal and 15.6 mm rostral to the interaural plane, the horizontal planes situated between 16.5 and 22.5 mm from the brain’s dorsal surface, and the sagittal planes between 0 and 6 mm from the medial wall. The coordinates correspond to area 29/31 in Paxinos et al. (2008, Wang et al., 2020; Foster et al., 2023).

We defined **VS** as lying within the coronal planes situated between 20.66 and 28.02 mm rostral to the interaural plane, the horizontal planes situated between 0 and 8.01 mm from the ventral surface of the striatum, and the sagittal planes between 0 and 8.69 mm from the medial wall. Note that our recording sites were targeted towards the nucleus accumbens core region of the VS. This dataset was used in Strait et al. (2015 and 2016).

We confirmed the recording location before each recording session using our Brainsight system with structural magnetic resonance images taken before the experiment. Neuroimaging was performed at the Rochester Center for Brain Imaging on a Siemens 3T MAGNETOM Trio Tim using 0.5 mm voxels or in the Center for Magnetic Resonance Research at UMN. We confirmed recording locations by listening for characteristic sounds of white and gray matter during recording, which in all cases matched the loci indicated by the Brainsight system.

### Electrophysiological techniques and processing

Either single (FHC) or multi-contact electrodes (V-Probe, Plexon) were lowered using a microdrive (NAN Instruments) until waveforms between one and three neuron(s) were isolated. Individual action potentials were isolated on a Plexon system (Plexon, Dallas, TX) or Ripple Neuro (Salt Lake City, UT). Neurons were selected for study solely on the basis of the quality of isolation; we never preselected based on task-related response properties. All collected neurons for which we managed to obtain at least 300 trials were analyzed; no neurons that surpassed our isolation criteria were excluded from analysis.

### Eye-tracking and reward delivery

Eye position was sampled at 1,000 Hz by an infrared eye-monitoring camera system (SR Research). Stimuli were controlled by a computer running Matlab (Mathworks) with Psychtoolbox and Eyelink Toolbox. Visual stimuli were colored rectangles on a computer monitor placed 57 cm from the animal and centered on its eyes. A standard solenoid valve controlled the duration of juice delivery. Solenoid calibration was performed daily.

### Risky choice task

The task made use of vertical rectangles indicating reward amount and probability. We have shown in a variety of contexts that this method provides reliable communication of abstract concepts such as reward, probability, delay, and rule to monkeys (e.g. Azab et al., 2017 and 2018; Sleezer et al., 2016; Blanchard et al., 2014). The task presented two offers on each trial. A rectangle 300 pixels tall and 80 pixels wide represented each offer (11.35° of visual angle tall and 4.08° of visual angle wide). Two parameters defined gamble offers, *stakes* and *probability*. Each gamble rectangle was divided into two portions, one red and the other either gray, blue, or green. The size of the color portions signified the probability of winning a small (125 μl, gray), medium (165 μl, blue), or large reward (240 μl, green), respectively. We used a uniform distribution between 0 and 100% for probabilities. The size of the red portion indicated the probability of no reward. Offer types were selected at random with a 43.75% probability of blue (medium magnitude) gamble, a 43.75% probability of green (high magnitude) gambles, and a 12.5% probability of gray options (safe offers). All safe offers were excluded from the analyses described here, although we confirmed that the results are the same if these trials are included. Previous training history for these subjects included several saccade-based laboratory tasks, including a cognitive control task (Hayden et al., 2010), two stochastic choice tasks (Blanchard et al., 2014), a foraging task (Blanchard and Hayden, 2015), and a discounting task (Pearson et al., 2010).

On each trial, one offer appeared on the left side of the screen and the other appeared on the right. We randomized the sides of the first and second offer. Both offers appeared for 400 ms and were followed by a 600-ms blank period. After the offers were presented separately, a central fixation spot appeared, and the monkey fixated on it for 100 ms. Next, both offers appeared simultaneously and the animal indicated its choice by shifting gaze to its preferred offer and maintaining fixation on it for 200 ms. Failure to maintain gaze for 200 ms did not lead to the end of the trial but instead returned the monkey to a choice state; thus, monkeys were free to change their mind if they did so within 200 ms (although in our observations, they seldom did so). Following a successful 200-ms fixation, the gamble was resolved and the reward was delivered. We defined trials that took > 7 sec as inattentive trials and we did not include them in the analyses (this removed ∼1% of trials). Outcomes that yielded rewards were accompanied by a visual cue: a white circle in the center of the chosen offer. All trials were followed by an 800-ms intertrial interval with a blank screen.

### Choice behavior model

Previous analysis and modeling of this behavioral data indicate monkeys make choices with a subjective value estimate of a risk-seeking attitude towards offer size (stakes) and a probability estimate well approximated by a prelec function. Subjects are assumed to choose according to the difference in offer one and offer two subjective values (Δ). Here, we also consider the role of reward rate on modulating this logistic choice function as predicted by rational-inattention theories.

All choice model maximum-likelihood optimization was performed using Scipy.optimize in Python with a binary-cross entropy loss. Each model was fit on a per-session basis. Choices were fit with a logistic choice model simultaneously with the power-function for utility (stakes) weighting and the prelec function for probability. The subjective value term for each offer was created by multiplying the utility and probability terms. Model-selection was performed by fitting all variants of the models, and compared using Akaike Weights (Wagenmarker and Farrell, 2004). The full logistic choice model in log-linear form was designed as follows, with linear and interaction regressors:

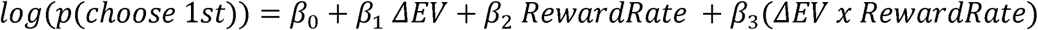

### Pupillometry

Blinks and missing data were cleaned from the pupil data by linearly interpolating the missing points. Pupil time-series were low-pass filtered at 15 Hz (butterworth, 2nd order), smoothed with a savitkgy-golay filter (window size 5 and poly order 3). The filtered time-series were then epoched to each offer window, starting at the fixation cross, up till the offer 2 memory window offset (2000 ms after the onset of 1st offer). The time-series were reference level corrected by subtracting the grand mean.

Pupil differences between low and high-reward average conditions were predicted to be different. This was assessed by computing the separate mean pupil trace for 33% and 66% percentiles of reward rate for each session. We computed the mean condition difference across subjects, and used a condition label permutation t-test (1000 permutes) at each time point. The true mean difference was compared against the permutations to establish a p-value.

### Neural Decoding

Decoding of either subjective value or reward rate was done using a linear support vector machine (SVM; LinearSVC in sklearn), with stratified 5-fold cross-validation, with a re-sampling of 90% of the maximum possible trials, repeated forty times. Decoding was performed using a 180 ms window with 40 ms moving window. All decoding analysis was performed using all subjects for a given brain area.

Reward rate decoding involved using the 33% and 66% percentiles to split the trials into a low and high reward rate, respectively. Because decoding analysis of subjective value also aimed to compare to value decoding at low-and high-reward rates, we conditioned the 33% and 66% percentiles of subjective value on the 33% and 66% percentiles of reward rate. Value decoders were then fit separately for each reward rate level. To compute decoder significance, for each condition, we also permuted the target labels and refit the decoder. This was performed 500 times. We considered decoding significant when *p<0.05* for at least two adjacent time-windows. For comparing decoding accuracy of value between reward rate conditions, we used the fitted decoders to correct labels. We permuted the accuracy scores between conditions, 1000 times to build a null distribution and compute a p-value at each time-point. Multiple comparisons across time-points were corrected using a false discovery rate.

### Cross-temporal decoding (CTD)

Dynamics of value coding was computed by using the time-point specific linear SVM computed above for value, in each of the 5 k-folds, and testing on all of the trials for the other time-point. The training time-point *i* was tested on time-point *j*, where the significance for CTD was determined using the permutation threshold determined using the permuted decoder for training on time-point *j*; this is the same threshold used for value decoding as described in *Neural Decoding*.

### Subspace orthogonality

The alignment or orthogonality of value coding reward rate specific subspaces was determined by computing a bootstrap correlation between value decoding (SVM) weights. The SVM weights define a one-dimensional axis in the neuron firing space that vary specifically with offer value. To compute the subspace orthogonality, we first averaged the weights across the 5 folds for each of the forty value decoding runs (subsampled trials). Averaging was done separately for both low and high reward rate decoder weights, yielding 80 total sets of weights.

We then computed the full set of correlations between these weights, correlating each low reward rate to each high reward rate set of weights. A noise threshold was computed to determine difference from zero by repeating the same procedure using the permuted sets of decoder weights. To determine whether subspaces were significantly semi-orthogonal (less than 1) we followed a previous procedure (Kimmel et al., 2020) and computed a ceiling threshold in two-steps. First, we compute all of the correlations between weights for each reward rate condition. This yields a separate vector of subspace correlations for the low and high reward rate conditions. Each of these correlations are then multiplied elementwise, and square root correcting, yielding a threshold distribution for testing of correlations significantly less than 1. Significance testing for either greater than noise or less than 1 (test of semi-orthogonality) was computed using a z-test that compared the mean actual subspace correlation to the distributions of noise and ceiling correlations.

